# Ultrasensitive allele inference from immune repertoire sequencing data with MiXCR

**DOI:** 10.1101/2023.10.10.561703

**Authors:** Artem Mikelov, George Nefediev, Alexander Tashkeev, Oscar L. Rodriguez, Diego A. Ortmans, Valeriia Skatova, Mark Izraelson, Alexey Davydov, Stanislav Poslavsky, Souad Rahmouni, Corey T. Watson, Dmitriy Chudakov, Scott D. Boyd, Dmitry Bolotin

## Abstract

Allelic variability in the adaptive immune receptor loci, which harbor the gene segments that encode B cell and T cell receptors (BCR/TCR), has been shown to be of critical importance for immune responses to pathogens and vaccines. In recent years, B cell and T cell receptor repertoire sequencing (Rep-Seq) has become widespread in immunology research making it the most readily available source of information about allelic diversity in immunoglobulin (IG) and T cell receptor (TR) loci in different populations. Here we present a novel algorithm for extra-sensitive and specific variable (V) and joining (J) gene allele inference and genotyping allowing reconstruction of individual high-quality gene segment libraries. The approach can be applied for inferring allelic variants from peripheral blood lymphocyte BCR and TCR repertoire sequencing data, including hypermutated isotype-switched BCR sequences, thus allowing high-throughput genotyping and novel allele discovery from a wide variety of existing datasets. The developed algorithm is a part of the MiXCR software (https://mixcr.com) and can be incorporated into any pipeline utilizing upstream processing with MiXCR.

We demonstrate the accuracy of this approach using Rep-Seq paired with long-read genomic sequencing data, comparing it to a widely used algorithm, TIgGER. We applied the algorithm to a large set of IG heavy chain (IGH) Rep-Seq data from 450 donors of ancestrally diverse population groups, and to the largest reported full-length TCR alpha and beta chain (TRA; TRB) Rep-Seq dataset, representing 134 individuals. This allowed us to assess the genetic diversity of genes within the IGH, TRA and TRB loci in different populations and demonstrate the connection between antibody repertoire gene usage and the number of allelic variants present in the population. Finally we established a database of allelic variants of V and J genes inferred from Rep-Seq data and their population frequencies with free public access at https://vdj.online.

## Introduction

Adaptive immune repertoire diversity plays a crucial role in shaping the immune response and forming immunological memory. Most immune repertoire research has focused primarily on somatically derived immune receptor diversity, namely V(D)J recombination and somatic hypermutation (SHM) diversity. In recent years, however, the extent of population diversity has begun to be appreciated at both the immunoglobulin (IG) (Gidoni et al., 2019; Mikocziova et al. 2021; Corcoran et al. 2023; Rodriguez et al. 2023; Gibson et al. 2022) and T cell receptor (TR) loci (Omer et al., 2022; M. Corcoran et al., 2023; Rodriguez et al. 2022). The functional significance of allelic variation in adaptive immune loci has also been recognized in the context of influenza, HIV and COVID-19 immunity and vaccination (Avnir et al. 2016; Lee et al. 2021; Leggat et al. 2022; Pushparaj et al., 2022).

Rep-Seq experiments sequencing repertoires of adaptive immune receptors encoded by recombined germline V, D and J genes have become a major source of information about adaptive immune functions in health and disease. In recent years, Rep-Seq has been utilized to discover many novel alleles in TR and IG loci, becoming one of the major sources of information of the allelic diversity of TR and IG genes in different populations. However, the major obstacle for utilizing Rep-Seq datasets for genotyping and allelic discovery is the presence of somatically hypermutated sequences in most available immunoglobulin Rep-Seq datasets, along with the PCR and sequencing errors which affect both BCR and TCR repertoire datasets. Hot-spot hypermutations and sequence errors have significantly hindered the ability to clearly detect individual polymorphisms. We aimed to overcome these issues with the algorithm described in this paper. However, there are two other challenging obstacles for accurate genotyping and haplotyping of TR and IG loci using Rep-Seq data only. Common structural variants (SVs) in IG loci (Rodriguez et al. 2023), especially gene duplications, in some cases make it hard to unequivocally map a sequence fromRep-Seq data to a particular germline gene without an additional source of information. Further, some alleles exhibit low usage levels, precluding their detection with Rep-Seq. Despite these limitations, Rep-Seq data remain valuable for applications focused on functional adaptive immune repertoires and their fluctuations in different conditions.

The ability to precisely call known allelic variants and infer novel ones from the same Rep-Seq data could enable new analyses of germline variation contribution to immune responses, and also improve the accuracy of many existing downstream approaches. There are several published methods for genotyping and allelic inference of V and J genes from Rep-Seq data (Table 1, numbers 2-5), however, each has important limitations. TIgGER (Gadala-Maria, Yaari, Uduman, & Kleinstein, 2015; Gadala-Maria et al., 2019) and Partis (Ralph & Matsen, 2019) are based on the idea that allelic sequence variants show a distinctive pattern over the background of SHMs, making SHMs presence in the dataset essential for reliable inference. This is not the case for many datasets, most obviously for T-cell receptor repertoire data and naive B-cell repertoires. On the other hand, IgDiscover (M. M. Corcoran et al., 2016), a very robust and reliable tool for novel allele inference, requires data without hypermutations, thus excluding much published immunoglobulin repertoire data.

**Table 1.**
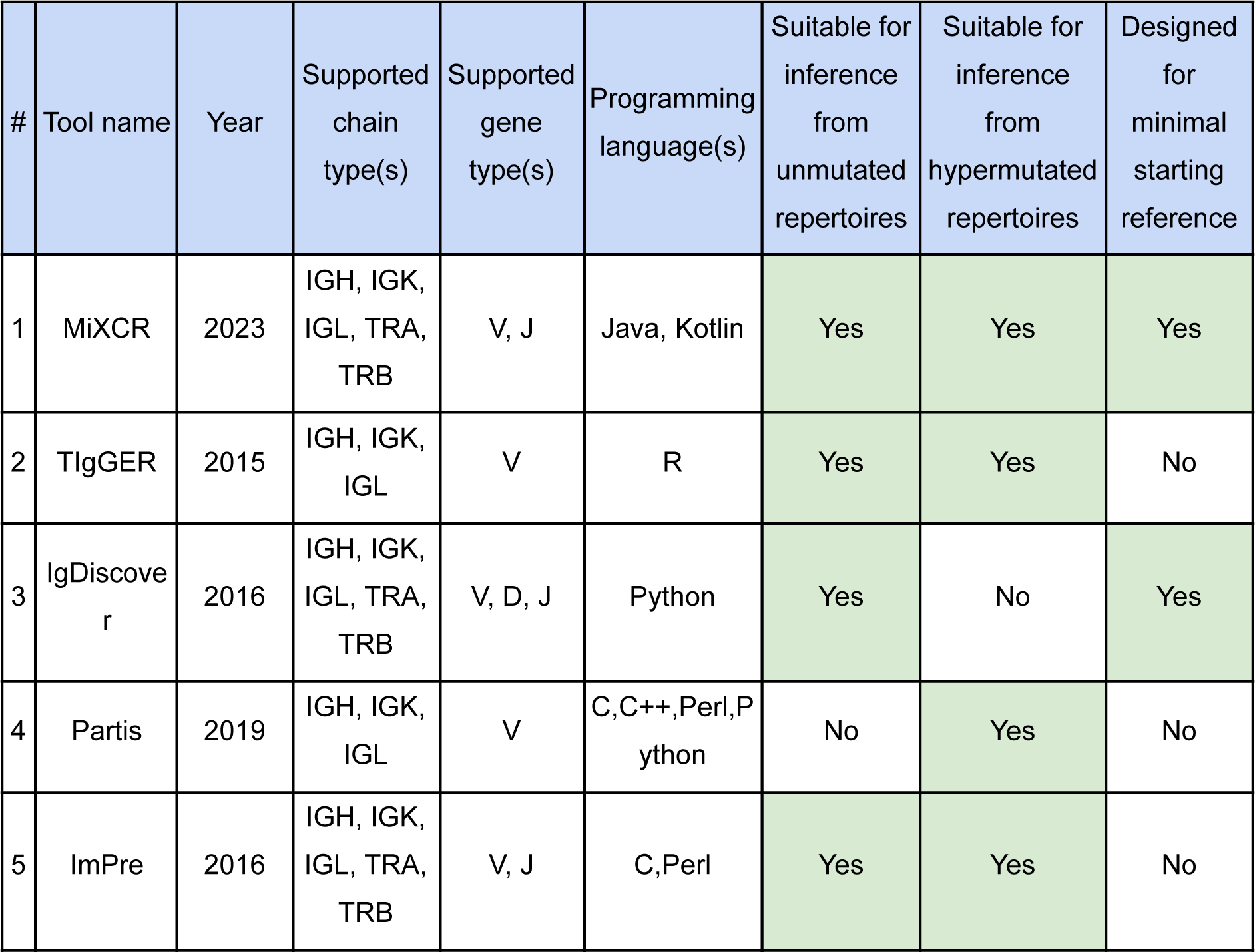
Tools for novel allele variants inference from Rep-Seq data and their characteristics.

Existing tools also require considerable depth of Rep-Seq data for reliable allele inference (for example, IgDiscover recommends at least 750,000 sequencing reads per individual library). Such sequencing depth is costly and not available for most publicly available Rep-Seq datasets. Here we present an algorithm for allelic inference and genotyping from both hypermutated and non-hypermutated repertoires, with low sequencing depth requirements. The algorithm performs well starting with a minimalistic gene reference library of only one allele for each gene, and even with some genes missing. These features make the tool especially useful for studying allelic diversity in non-model species where reference gene libraries are sparse and incomplete. The developed approach is integrated in MiXCR software and is available as the `findAlleles` command.

The International ImMunoGeneTics Information System (IMGT®), established in 1989, is the oldest widely available source of information about immune receptors, including alleles. Recent advancements in high-throughput adaptive immune receptor repertoire sequencing (Rep-Seq) methods enabled a broader view of alleles, and many tools were developed to infer allelic variants from such data. In 2017, the AIRR-Community (a network of over 300 practitioners in the field of Rep-Seq, www.airr-community.org) and IMGT® agreed on a process for adding new alleles inferred from Rep-Seq data to the IMGT® database (Ohlin et al., 2019). The AIRR Community also introduced the Open Germline Receptor Database (OGRDB, https://ogrdb.airr-community.org/, Lees et al., 2020), to track the addition of new alleles, but this currently harbors a limited number of sequences (e.g. 41 for human IG heavy and light chains). Although being the most recognized source of germline immunoglobulin sequence data, IMGT® lacks information on population allele frequencies and harbors sequences ‘mapped’ to the identified genes at the specific genomic locations. However, structural variation is quite common (Rodriguez et al. 2023) in the IG and TR loci, while most new IG and TR sequence data is coming from Rep-Seq experiments, and can be hard to map to a particular gemline locus position. Other databases of immune receptor gene alleles have been introduced, such as pmTR (Dekker, van Dongen, Reinders, & Khatri, 2022, https://pmtrig.lumc.nl/), IgPdb (https://cgi.cse.unsw.edu.au/∼ihmmune/IgPdb) and Karolinska Institutet human T-cell receptor database (M. Corcoran et al., 2023, https://gkhlab.gitlab.io/tcr/). A comprehensive and well-maintained database of immune receptor gene alleles, including allelic variants inferred from Rep-Seq, is VDJbase (Omer et al. 2020, https://vdjbase.org/). However, VDJbase is not seamlessly integrated with any of the analysis tools, and using it for Rep-Seq data analysis requires conversion of sequence data formats. To accompany the MiXCR software, we have developed VDJ.online (https://vdj.online/library), a free and open database of immune receptor allelic sequences that enables examination, comparison, and downloading of sequences.The VDJ.online reference library is supplied with the MiXCR, allowing seamless Rep-Seq data processing with accurate V and J gene annotation, genotyping, and novel allele inference.

## Results

### Novel approach to V and J gene allele variants inference and genotyping

The main challenge of allelic inference from Rep-Seq data is the presence of hypermutations, PCR and sequencing errors, with a large fraction of them being hot-spot mutations occurring simultaneously in unrelated clones. We have overcome this challenge by consecutively applying several filters based on two major measures. The first one is the lower diversity bound, estimated as the number of unique combinations of J and V genes and CDR3-lengths of clonotypes. The second measure is based on the number of clonotypes with unmutated J and V genes. Filters are applied both at individual mutation and at mutation set levels (see Methods for the detailed description). The germline-encoded mutations in CDR3 are recovered, when possible, by using non-mutated clonotypes matching the inferred variants in the rest of the sequence. This approach allows both to infer novel (undocumented) V and J gene alleles and to perform genotyping with high sensitivity and precision.

### Benchmarking of the V and J gene allele variants inference and genotyping

To assess the performance of the developed algorithm we utilized publicly available datasets (Rodriguez et al. 2023) containing both Rep-Seq data and highly reliable genotyping data of the IGH locus reconstructed using Pacific Biosciences HiFi long-read sequencing. For the sake of comparison we utilized 33 Rep-Seq data sets of sufficient sequencing depth (> 500,000 sequencing reads) and at least 3,000 unique full-length clonotypes. We limited our comparison to tools which were suitable for this type of data, i.e. excluding IgDiscover, which requires unmutated datasets for the correct performance. As mentioned above ImPre is no longer supported, while the run time of Partis exceeded two weeks on a high-performance cluster, which could limit real-world applications. Thus, for the comparison we utilized TIgGER, which is also the most widely cited tool for this task. Upstream analysis, including sequence alignment to reference V and J gene libraries and defining the full-length clonotypes, was performed using the tools’ recommended pipeline, MiXCR’s `analyze` module (Bolotin et al., 2015, https://mixcr.com/mixcr/reference/mixcr-analyze/), and Presto (Vander Heiden et al., 2014) and Change-O (Gupta et al., 2015) from Immcantation framework (https://immcantation.readthedocs.io). For further details please see the Methods section. For the alignment step and V and J gene annotation we used a custom minimalistic gene set library with only one allelic variant per V and J gene, derived from a custom public genome reference to match the one used for the long-read based genotyping (Rodriguez et al. 2020). Then we performed allele variant inference and genotyping with both tools for all datasets containing more than 3,000 unique full-length clonotypes and compared the resulting individualized V and J gene libraries with the accurate genotype inferred with the next generation long-read sequencing (Rodriguez et al. 2023), comparing nucleotide sequences of the genes. We also excluded poorly expressed allelic variants as determined by aligning the reads to the individualized gene reference libraries. Thus, in our benchmarking we focused on the question of detection of particular V or J gene allele sequences in the participants’ Rep-Seq IGH data for the subsequent accurate clonotype annotation, which is crucial for many downstream applications of such data (e.g. lineage trees analysis). Importantly, we compared the abilities of both approaches using the sparse reference libraries and Rep-Seq data, containing varying amounts of errors and sequencing noise, including datasets incorporating unique molecular identifiers and not. Therefore, we consider our benchmarking relevant to real world applications where the data quality is typically far from ideal in many aspects.

MiXCR on average detected 98% of the allelic variants of the V genes supported by the long-read based genotyping, while TIgGER detected 81% of the V gene alleles (**Fig. 1A**). MiXCR produced on average 1 allele call not supported by long-read based genotyping, while TIgGER yielded 2 potential false positive calls (**Fig. 1B**). Interestingly, the recall of TIgGER improved remarkably (up to 94% on average) when the upstream analysis and allele inference was performed utilizing the full built-in reference library containing all of the known alleles **(Supplemental Fig. S1A**). However, the number of the allele calls not supported by the long-read sequencing also increased, up to 5 potential false-positive calls on average. **(Supplemental Fig. S1B**). For MiXCR, transition to full reference library resulted in only minor changes in performance **(Supplemental Fig. S1A, S1B**).

**Figure 1.**
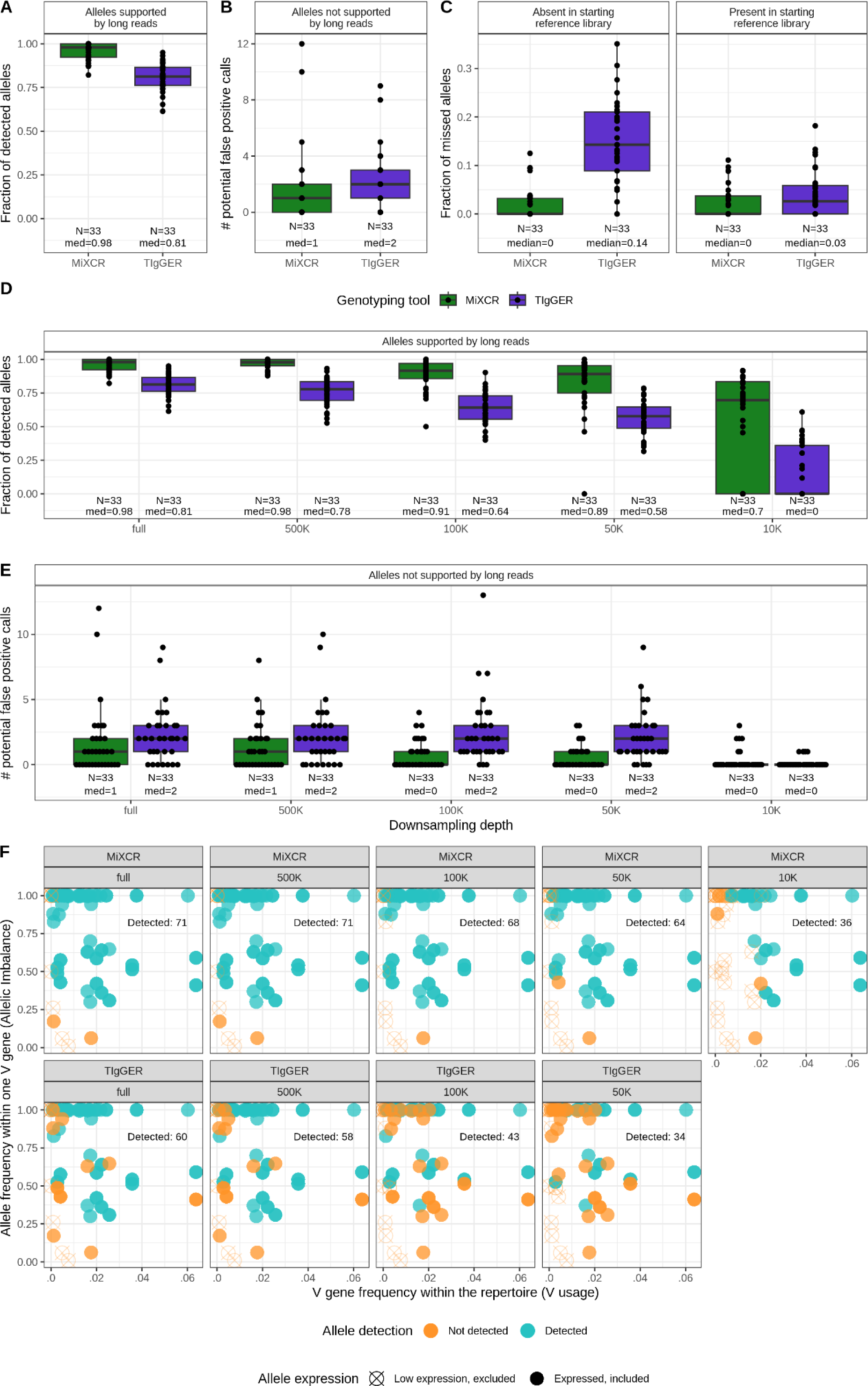
Detection of allelic variants of V genes by inference tools. a, Fraction of allele calls supported by long-read based genotyping. b, Number of allele calls not supported by long-read based genotyping. c, Fraction of alleles, missed by MiXCR or TIgGER, by presence in the initial reference library. d, e Sensitivity and specificity testing by downsampling each sample in the benchmarking dataset by 500,000, 100,000, 50,000 or 10,000 reads. d, fraction of identified allele calls supported by long-read based genotyping. e, number of identified allele calls not supported by long-read based genotyping. f, detection of the allele variants of V genes depending on V usage and allelic imbalance. Each dot represents a V gene allele present in the donor’s genotype confirmed by long-read sequencing. The upper row represents detection by the developed algorithm, the lower, allele detection by the gold standard tool TIgGER. Columns represent different depths of downsampling by number of aligned reads, from right to left: full set of reads, 500,000, 100,000, 50,000, 10,000. V gene, and allele frequencies for each facet were calculated using the full set of reads and allele-resolved V and J gene reference library. Alleles excluded due to low expression (<10 clonotypes), are represented as empty crossed points.

The difference in the number of called alleles between the two algorithms was also apparent when we compared rates of detection of the *de novo* inferred alleles. TIgGER did not detect on average 14% of alleles absent in the starting reference gene library, while MiXCR missed none of the alleles (**Fig. 1C**).

To test the sensitivity of the approaches we also downsampled the dataset to 500,000, 100,000, 50,000 and 10,000 raw sequencing reads. MiXCR allele detection rates decreased by 9 percentage points down to 89% on average when downsampled to 50,000 reads, which is more than 10x downsampling for all of the datasets. TIgGER detection rates also deteriorated by 23 percentage points, detecting on average 58% of alleles with 50,000 reads. At the extreme level of downsampling by 10,000 sequencing reads MiXCR was able to detect 70% of alleles on average, while TIgGER yielded an error for 21 of the samples due to the low number of clones assigned to any of the V genes (**Fig. 1C**). For MiXCR, the detection of the alleles clearly depended on the two variables - the frequency of the V gene in a particular repertoire and the imbalance in usage between different alleles for a particular V gene. For TIgGER, these parameters appeared to have little influence on detection rates (**Fig. 1D**).

For the task of detecting J gene allelic variants, for which could not be performed with TIgGER, MiXCR yielded 100% sensitivity and specificity even with the datasets downsampled to 50,000 reads (**Figure S2**).

### Numerous IGH, TRA and TRB novel alleles detected using MiXCR allele inference

To investigate allelic diversity in human populations we applied the developed algorithm to a large collections of IGH (450 individuals) and full-length TRA and TRB (134 individuals) Rep-Seq datasets. The MiXCR allele inference and genotyping pipeline resulted in identification of both known and previously undocumented alleles, 384 IGHV, 128 TRAV, 144 TRBV, 14 IGHJ, 64 TRAJ, and 14 TRBJ in total.

Numerous previously undocumented alleles, absent from major databases mentioned above, were detected: 183 IGHV, 33 TRAV, 7 TRAJ, 41 TRBV (**Fig. 2A-D, F**). Of note, we did not detect any novel variant for any of the IGHJ and TRBJ genes (**Fig. 2E**). All of the novel alleles sequences were contributed to the public database of allelic variants and are available for download at https://vdj.online/library (**Supplemental Fig. S3**).

**Figure 2.**
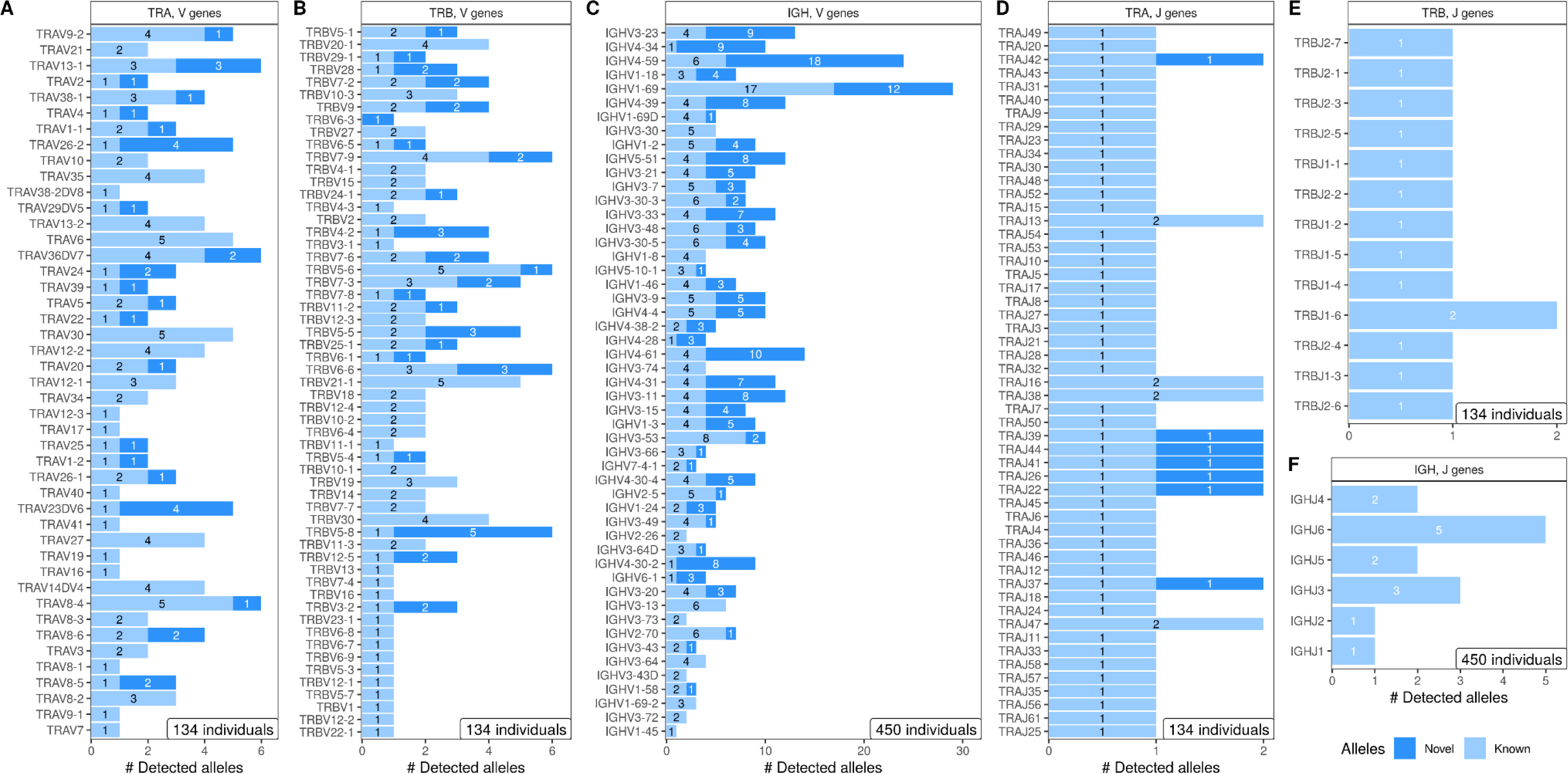
Number of observed novel and known alleles. a, TRAV. b, TRBV. c, IGHV. d, TRAJ. e, TRBJ. f, IGHJ

### Divergent allele frequency distribution in IGHV genes in African population

The considered IGH Rep-Seq datasets included repertoires from African, Asian, Caucasian and Hispanic/Latino individuals (Fig. 3A), allowing us to investigate differences in IGHV and IGHJ allelic distributions between these major population groups. We did not observe significant differences in the total number of detected novel IGHV and IGHJ alleles between those groups (**Fig. 3B**). The number of detected alleles was similar for most of the V genes with the exception of IGHV5-51, IGHV1-69 and IGHV4-30-2 (**Fig. 3D**).

**Figure 3.**
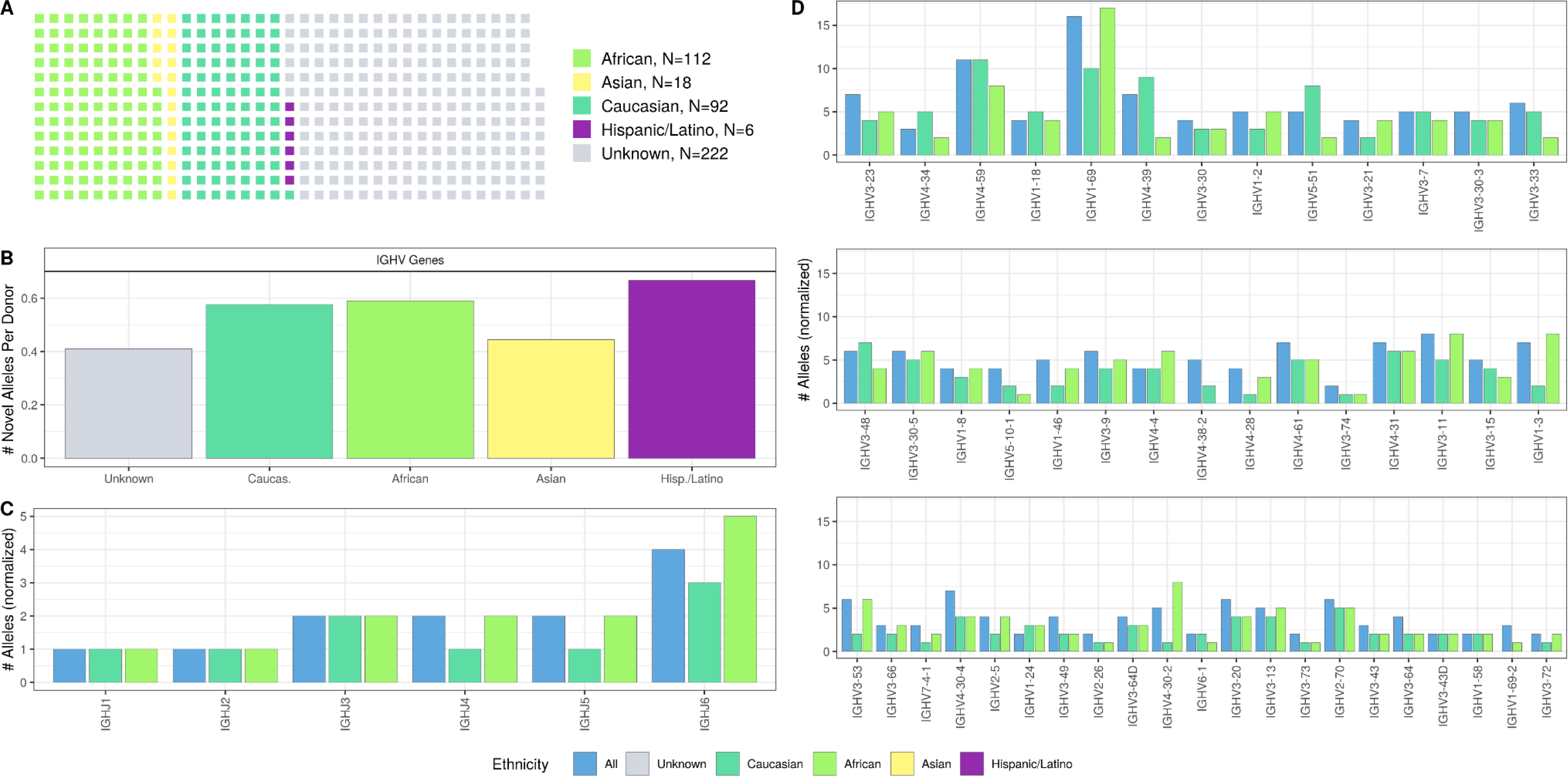
IGHV and IGHJ allelic diversity by major ethnic groups. a, Cohort composition. b, Number of detected novel alleles, normalized per number of individuals. c, Total number of detected alleles by IGHJ gene in caucasian, african and general population, normalized by downsampling to a fixed number of individuals (N=92). d, Total number of detected alleles by IGHV gene in caucasian, african and general population, normalized by downsampling to a fixed number of individuals (N=92)

On the other hand, the allele frequency distribution in the African population was different than that of other populations for 19 IGHV genes (**Fig. 4, Supplemental Fig. S4**). The same difference was also observed for two of the IGHJ genes: IGHJ3 and IGH6 (**Supplemental Fig. S5**). In TRBV and TRAV loci for most of the gene frequencies, distributions were heavily skewed towards particular single allele variants (**Supplemental Fig. S6, S7**), which may be attributed to a more homogeneous cohort composition by ethnicity, with the predominant majority of participants being of European descent.

**Figure 4.**
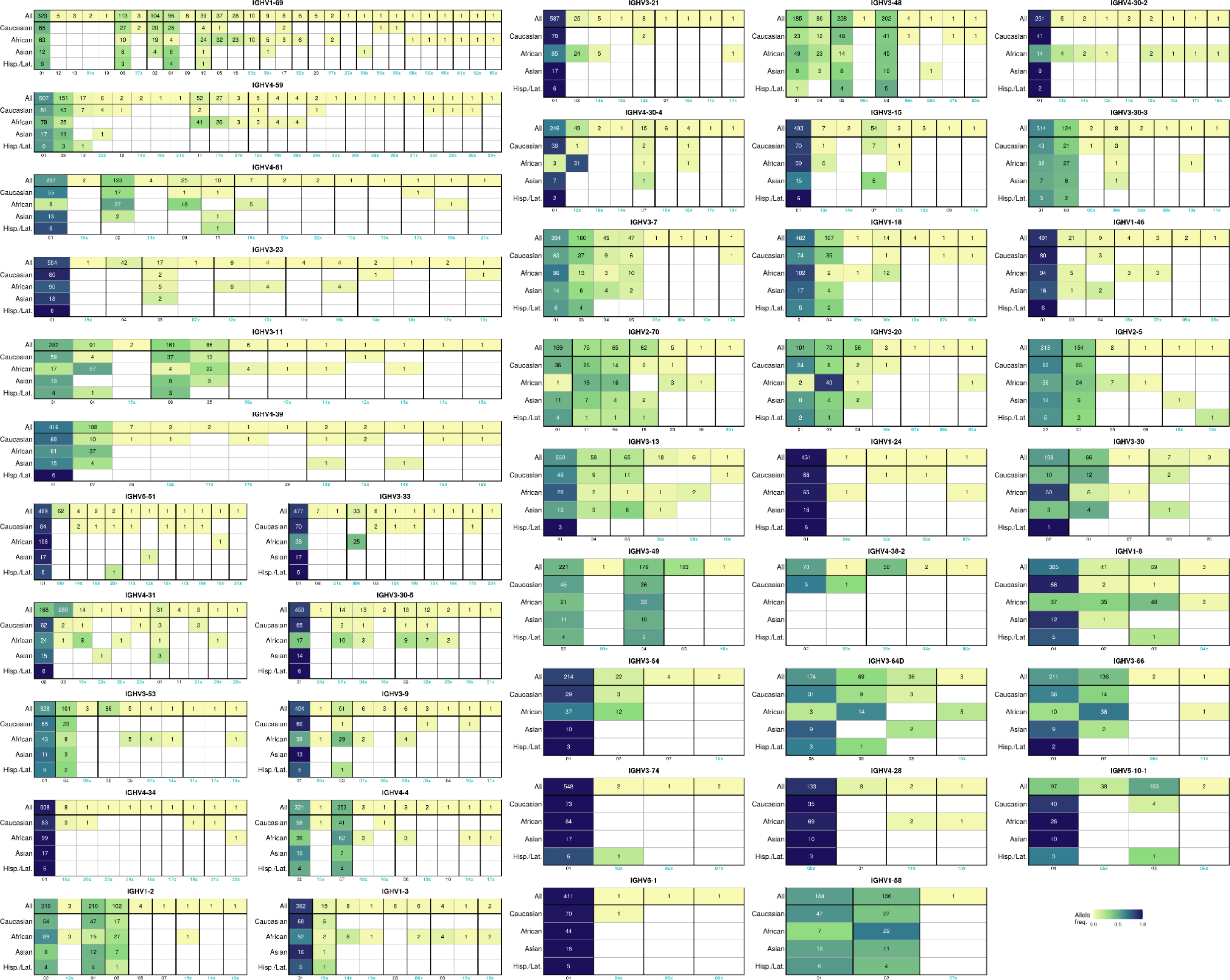
IGHV gene allele frequencies in major ethnic groups. Each column in heatmaps represents a particular allele; numbers for novel alleles, are colored in green, and concatenated with letter ‘x’; bold lines separate groups of alleles with different amino acid sequences; groups of alleles with the same amino acid sequence are ordered by the aggregated frequencies of alleles; alleles within groups are order by allele frequency in the general population. Color represents the allele frequency within the ethnic group; numbers in cells represent the number of occurrences of the corresponding allele.

## Discussion

Immune receptor repertoire sequencing datasets have become a valuable source of information for studying immune responses across different health conditions, tissues and cell subsets. Recently developed specialized algorithms (Gadala-Maria et al., 2019; M. M. Corcoran et al., 2016; Zhang et al., 2016; Ralph & Matsen, 2019) allow inference of allelic variants of V and J genes of adaptive immune receptors from Rep-Seq data, scaling up the process of novel allele discovery and allowing Rep-Seq data analysis using individualized gene reference libraries, which significantly increases the accuracy and quality of many types of downstream repertoire analyses. However, all of the current approaches demand high sequencing depth and a significant number of unique receptor sequences for the analysis. Moreover, many prior approaches do not allow allelic inference from both hypermutated and non-hypermutated repertoires. The most comprehensive approach for precise genotyping and allelic inference, utilizing long-read sequencing of the immune receptor gene loci (Gibson et al., 2022; Rodriguez et al. 2023; Rodriguez et al., 2020), has the greatest accuracy, and also allows for the investigation of structural variants. Although being the most desirable and accurate way to obtain donor-specific V and J gene genotypes, this methodology is costly and requires special experimental procedures. Here, we addressed this unmet need by developing an alternative approach for inferring allelic variants of V and J genes directly from Rep-Seq data, offering improved sensitivity and accuracy compared to existing tools.

Our method allows for successful allelic inference from datasets downsampled to as few as 50,000 sequencing reads. Moreover, the algorithm applicability is not restricted to a particular type of Rep-Seq data; it can be applied to both repertoires containing hypermutated sequences (e.g., IGH repertoires generated from any isotype) as well as datasets containing only non-hypermutated sequences (e.g. TCR-repertoires). Multiple filtering steps integrated into our pipeline prevent false-positive polymorphism calling which typically arises due to the presence of hot-spot hypermutations and PCR and sequencing errors. Furthermore, we demonstrate high sensitivity and specificity of the approach utilizing a very sparse starting reference gene library, containing only one allelic variant per gene, which makes it even more useful for studying allelic diversity in non-model species for which V and J gene reference libraries are incomplete and lack allelic variants. The developed approach is integrated within the MiXCR (Bolotin et al., 2015, https://mixcr.com) pipeline for immune-repertoire analysis, and allows seamless allelic inference and re-aligning repertoires to a personalized reference library.

Applying the developed approach to large collections of IGH, TRA and TRB repertoire datasets, we were able to identify a large number of previously undocumented V and J gene alleles. This allowed us to establish a database of allelic variants integrated and MiXCR and publicly available at VDJ.online database (https://vdj.online/library).

Differences in allele frequency distributions may have major implications for susceptibility of different populations to diseases and vaccination outcomes (Avnir et al., 2016). Large sample sizes (450 individuals for IGH and 134 for TRA/TRB) allowed us to estimate allele frequencies for most of the studied genes in the population. For IGHV and J gene allelic variants we identify striking differences in allele frequency distributions between African donors and other major population groups. We also contributed the information on V and J gene allele frequencies to VDJ.online, making it a valuable public resource of such information. Having incorporated this database of allelic variants into the MiXCR platform, we hope that it will facilitate further advancement in the immune repertoire analysis field, adding the dimension of allele analysis with little additional effort and cost to many further studies.

## Materials and methods

### Allele variants detection algorithm

The algorithm utilizes alignment and clonotype assembly information from the upstream Rep-Seq data processing, specifically mutation calls from reference V and J gene reference library for BCR or TCR clonotypes and V and J gene annotations, readily available after running the ‘analyze’ command in the MiXCR software (Bolotin et al. 2015). The clonotype definition for the purpose of allele inference may vary depending on the region covered by sequencing.

Using these defined sets of mutations which differentiate the particular clonotype sequences from the corresponding reference V or J gene, the algorithm then separately infers alleles for V and J genes. For simplicity, we describe the algorithm steps for V genes only, the J gene inference follows the same logic:

1. Clonotypes are grouped by the V genes.
2. For each mutation within the group, including insertions and deletions, we define a set of clonotypes which contain this mutation.
3. The mutations are filtered based on the lower diversity bound, estimated as the number of unique combinations of J genes and CDR3-lengths of clonotypes containing that mutation. The mutations that don’t exceed a predefined threshold for the value are removed from each of the clonotype’s mutation sets.
4. Clonotypes are grouped by filtered mutation sets, including “empty” mutation sets, containing no mutations. The lower diversity bound is calculated for each of the groups as described above. Additionally, the number of clonotypes containing no mutations in J gene after filtering as described in step 3 is calculated. Mutation sets are then filtered by thresholds of these two parameters, resulting in a list of allele candidates.
5. Clonotypes are then assigned to the closest allele candidates. Clonotypes which can not be unambiguously assigned are filtered out. Lower bound of naive diversity is calculated as the number of unique combinations of J genes and CDR3-lengths for clonotypes with unmutated J gene sequences. Candidates are sorted by the score which represents the weighted sum of the lower bound of diversity and lower bound of naive diversity, calculated as described above. Formula for the score:

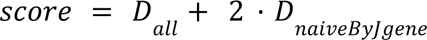 Where *D_all_* is the lower bound of diversity for all clonotypes; *D_naiveByJgene_* - lower bound diversity, calculated only for clonotypes with no mutations in J gene.
6. Candidates with the score not lower than 0.35 of the maximum score are then selected for the subject-specific gene set library.
7. The germline-encoded mutations in CDR3 are recovered using the non-mutated clonotypes, which totally match the inferred variants by the rest of the sequence excluding CDR3. Each position is considered if it has at least 5 clonotypes covering it and 70% nucleotide concordance. The right-most position in CDR3, which meets these criteria, is reported by MiXCR (`reliableRegion` field in tabular output). The rest of CDR3 is picked from the closest allele in the database.

The process for inferring J gene alleles is the same, however the initial grouping is performed by J genes and V genes are used for all of the filtering steps.

This stepwise approach based sequential filtering first on the level of individual mutation and then on the level of mutation sets dramatically reduces noise introduced by SHM and sequencing and PCR-errors. The threshold of 0.35 for the final allele filtering was initially chosen from a theoretical consideration of possible distributions of expressed alleles for a V gene allowing the presence of three allelic variants due to possible V gene duplications. This was then corroborated by examining empirical score distributions for alleles in sequencing of IGH repertoire of a healthy donor with known genotype; in this case the donor was different from the one in the benchmarking of the algorithm.

In case of a significant difference between the reference library and a particular individual’s genotype, the algorithm repeats the steps described above twice, with two different sets of parameters. The first step generates preliminary allele calls, which allows more precise estimation of numbers of clonotypes with unmutated gene sequences.

The algorithm always utilizes only one allele variant per gene as starting reference, preventing potential biases towards particular known sequences. In case there is a weak signal in a particular gene (usually represented by less than 20 clonotypes), the algorithm falls back to assigning one of the known alleles.

Finally, for all of the allele calls, the allele names are looked up in a reference database (the same as available at https://vdj.online/library) by exact match of nucleotide sequence. If there is no match the new name is derived from concatenation of the closest allele and sequence hash.

The described algorithm is integrated into MiXCR as the `findAlleles` command.

### Data collection and repertoire sequencing

For the benchmarking purposes we utilized IGH repertoire sequencing data, accompanied by a targeted long-read IGH locus sequencing from Rodriguez et al. 2023, selecting samples which had at least 500,000 sequencing (N=33), which was necessary for compatibility in downsampling experiments. IGH locus assembly and variant detection characterizing novel alleles were performed using iGenotyper (https://github.com/oscarlr/IGenotyper, Rodriguez et al. 2020) as previously described (Rodriguez et al. 2023).

For calculating population allele frequencies we used publicly available IGH Rep-Seq data from 6 published studies (total N=450) (Gidoni et al. 2019; Nielsen et al. 2020; Nielsen et al. 2019; Roskin et al. 2015; Davis et al. 2019; Rodriguez et al. 2022).

For generating high-quality full-length TCR repertoires, peripheral blood was collected from 134 individuals without major chronic immunological conditions at CHU of Liège, including COVID-19 patients and individuals after vaccination. 2.5 mL of blood was collected on PAXGene RNA tubes from each participant and stored at -80°C until use, RNA was extracted using the PAXgene Blood RNA Kit (Qiagen). cDNA libraries were generated using SMARTer Human TCR a/b Profiling Kit v2 (Takara Bio USA, San Jose, California, USA). Briefly, a rapid amplification of cDNA ends (RACE) approach with a template-switch effect was used to introduce 5’ adaptors during cDNA synthesis. cDNA corresponding to *TCRA* and *TCRB* transcripts was further amplified and prepared for sequencing, which was performed on a MiSeq instrument with paired-end 2×300 bp reads using the NovaSeq 6000 SP Reagent Kit v1.5 (500 cycles) (Illumina, San Diego, California, USA). The protocol was approved by the ethics committee of Liège University Hospital (approval numbers 2021-54 and 2020/107).

### Benchmarking of allele variants detection and genotyping

Processing of the Rep-Seq data was performed using MiXCR v4.4.0 (https://mixcr.com, Bolotin et al. 2015) upstream pipeline ‘analyze’ command, parallelized using GNU Parallel (Tange 2018). Importantly, for the alignment step and V and J gene annotation we used a custom minimalistic gene set library with only one allelic variant per V and J gene, derived from a custom public genome reference to match the one used for the long-read assembly (Rodriguez et al. 2023). After processing we excluded samples with the resulting number of full-length IGH clonotypes less than 3,000 (N=7), which probably related to samples either with low cell counts or with low RNA yield. Then the allelic variants were inferred and individual genotypes were reconstructed for each individual sample (N=33) with the algorithm described above integrated into MiXCR pipeline as `findAlleles` command.

To infer the alleles with the comparison tool, TIgGER (Gadala-Maria et al., 2019) we used the same set of samples (N=33). For initial Rep-Seq data processing we utilized tools pRESTO (Vander Heiden et al., 2014) and Change-O (Gupta et al., 2015), which are the part of the Immcantation framework along with TIgGER (https://immcantation.readthedocs.io), using commands and settings, recommended by the documentation. TIgGER v1.0.1 functions `findNovelAlleles` and `inferGenotype` were used for inferring novel alleles and reconstructing genotypes.

To test the sensitivity of the approaches we downsampled the dataset to 500,000, 100,000, 50,000 and 10,000 raw sequencing reads using seqtk (https://github.com/lh3/seqtk) v1.3 and applied the same upstream processing, allele inference and genotyping pipelines as for the full datasets.

The resulting sets of allele sequences were exported from both tools in fasta format and matched with the sequences of the alleles present in the genotype of the donor and the number of matches was determined. Importantly, due to the fact that IGH repertoire sequencing data utilized for comparison was derived using RNA-based technology, inference could be performed only for expressed V and J gene alleles. Thus, we excluded non-functional alleles and also those alleles from comparison which had less than 10 total clonotypes or less than 3 “naive” clonotypes with no mutation calls assigned to these alleles, when utilizing the same MiXCR v4.3.2 upstream pipeline, but with the individual allele-resolved V and J gene reference libraries constructed from long-read based genotypes. Also we have excluded from comparison genes which were not captured by the long-read sequencing. In particular, IGHJ genes were covered only for 9 of 33 considered individuals. For the benchmarking purposes, we excluded alleles for low abundance genes with too low abundance, as defined by each of the tools. TIgGER could not infer novel alleles for genes with less than 50 clonotypes assigned to it, so we excluded such alleles from comparison for TIgGER. MiXCR reports the low abundance genes for which the analysis is impossible with the parameters described above. The average number of clonotypes assigned to those genes was less than 10, we excluded such alleles from comparison for MiXCR too. Finally, we have not taken into account false negative and false positive polymorphism calls in the whole CDR3 region for TIgGER; for MiXCR we applied stricter criteria and have not considered false negative and false positive polymorphism calls outside `reliableRegion`, defined by the tool as described above.

### Novel allele inference and population frequencies

Processing of the Rep-Seq data for both BCR and TCR repertoires was performed using MiXCR v4.3.2 `analyze` command. Repertoires containing less than 3000 unique clonotypes were not used for downstream analysis. The algorithm described above for inferring novel alleles and genotyping was used by invoking `findAlleles` MiXCR command under default settings. The alleles lacking designated names by the International Union of Immunological Societies were labeled as undocumented.

For both TCR and immunoglobulin V and J gene alleles the number of haplotypes with these alleles were estimated using output tables from `findAlleles` command. Each case where the only one allele per gene was indicated was treated as a gene in homozygous state, thus not taking into account possible deletions of the genes on one of the chromosomes. We also limited our analysis with the genes detected in at least 15% of the donors. Allele frequencies were then calculated by dividing the number of haplotypes for a particular allele by the total number of haplotypes for this gene in the population. For the IGH data we also were able to calculate allele frequencies for four major ethnic groups - African, Asian, Caucasian, Hispanic/Latino (self-reported by participants, where missing assigned to “unknown”).

To evaluate pairwise similarity between IGH allele frequency distributions in different populations, we utilized Jensen-Shannon divergence, calculated using the following formula:

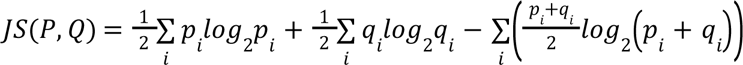

where P and Q represent the distributions of alleles of a particular V or J gene in two populations, and p_i_ and q_i_ represent frequencies of individual member *i* (one particular allele).

## Author Contributions

AM, GN, SP and DB conceived the study. GN and AM developed and benchmarked the described algorithm. OLR and CTW performed contig assembly and allele calling from long-read sequencing data. AM, DAO, MI, VS, AD, performed data analysis for the large population cohort. AT performed experimental work and SR supervised data collection for the TCR cohort. DC, SDB and DB supervised the study. AM wrote the manuscript with inputs from all authors.

