## Supplemental figures for "Ultrasensitive allele inference from immune repertoire sequencing data with MiXCR"

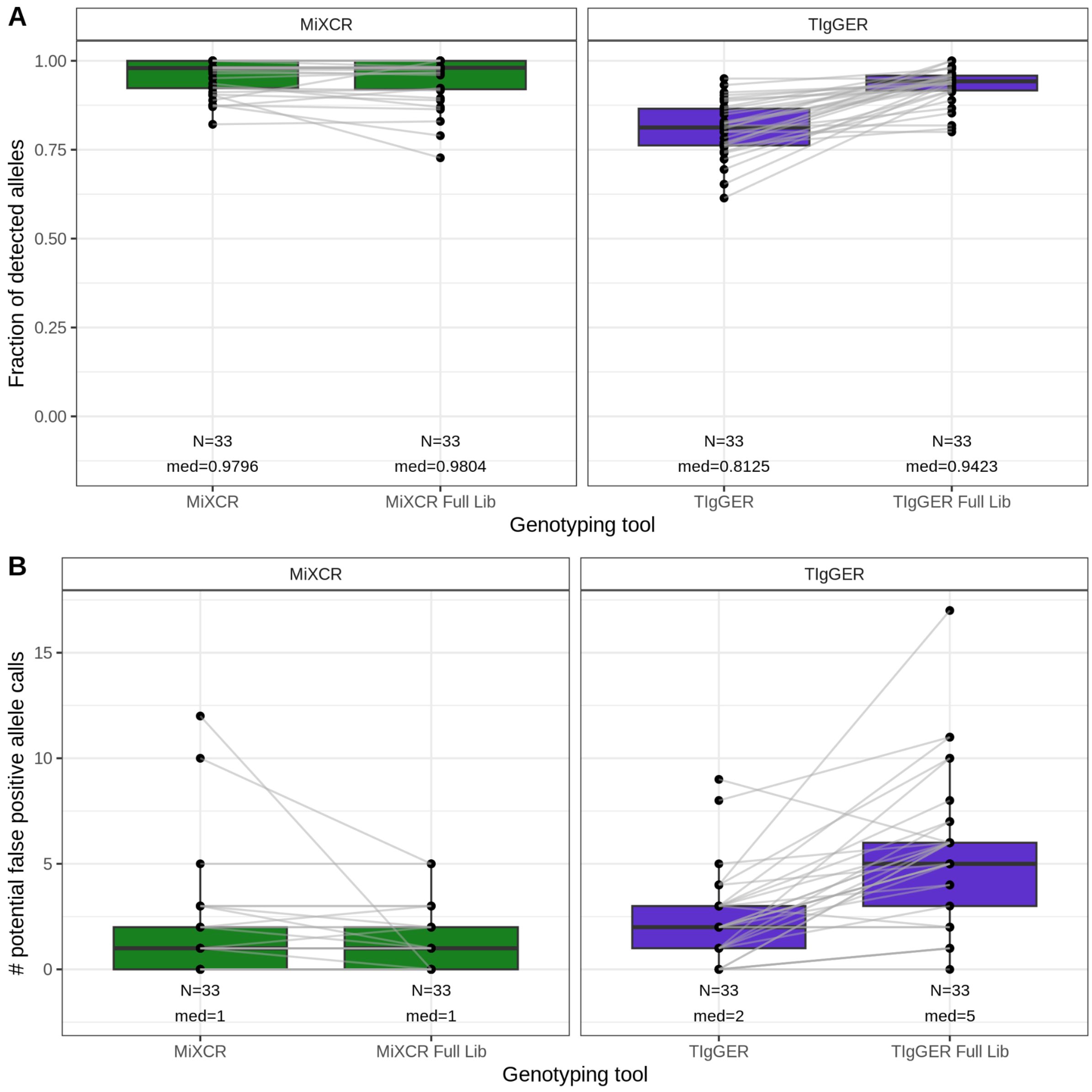


**Supplemental Figure S1.** Allele tool inference benchmarking using full reference libraries for each of the tools.

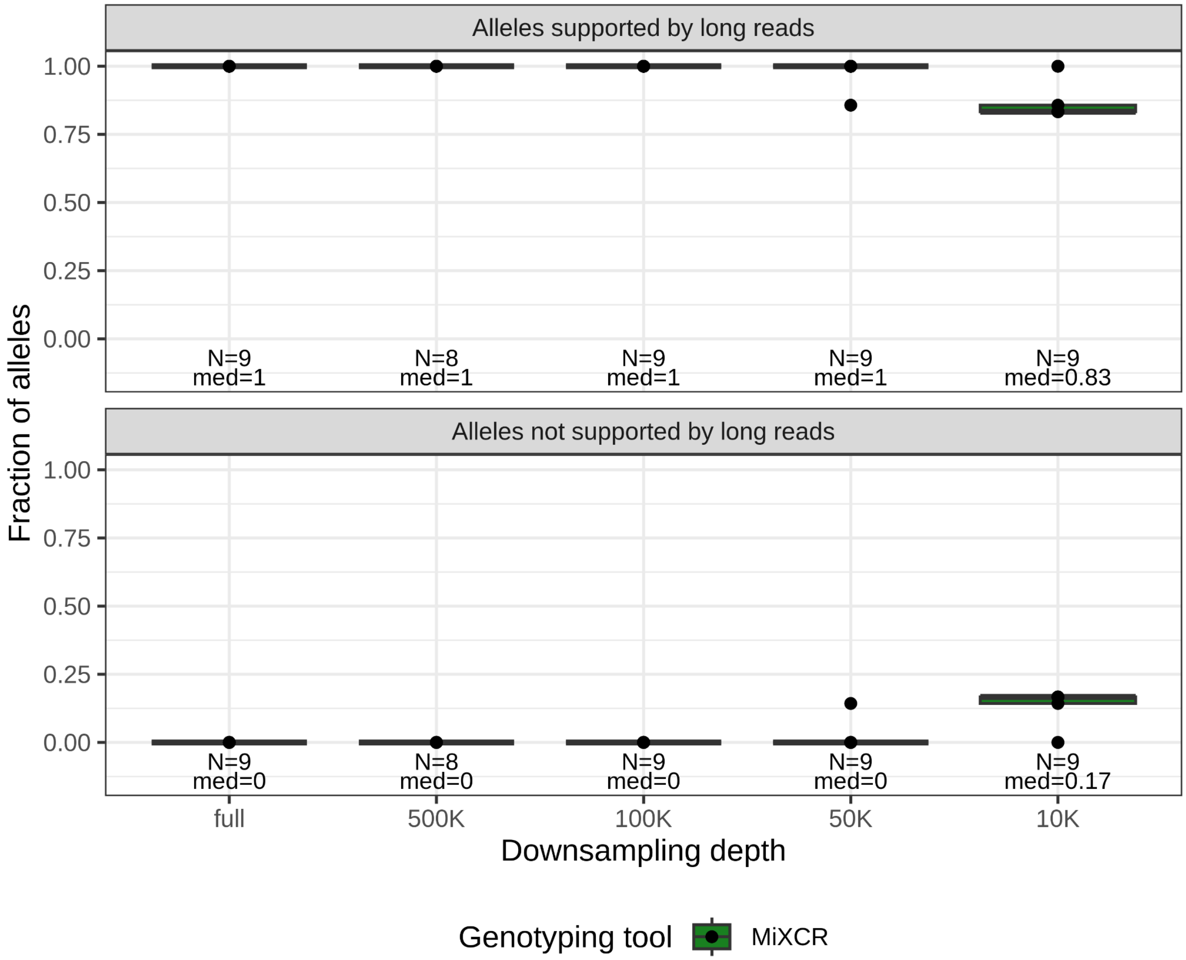


**Supplemental Figure S2.** Allele inference and genotyping of J gene allelic variants by MiXCR.

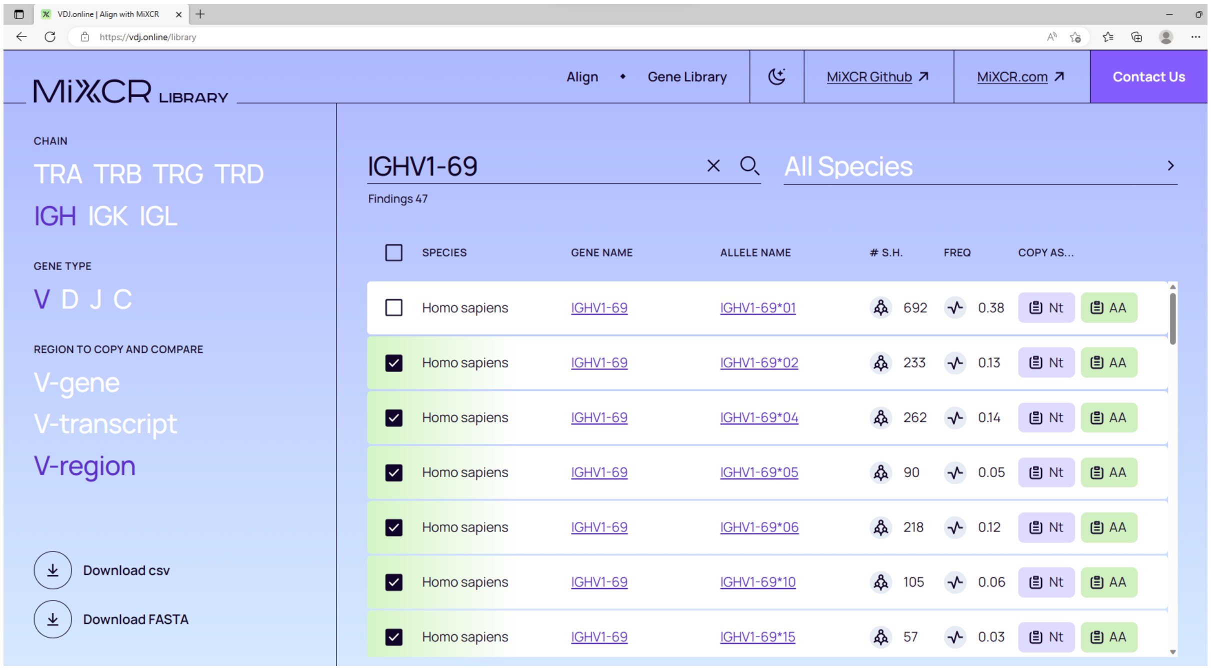


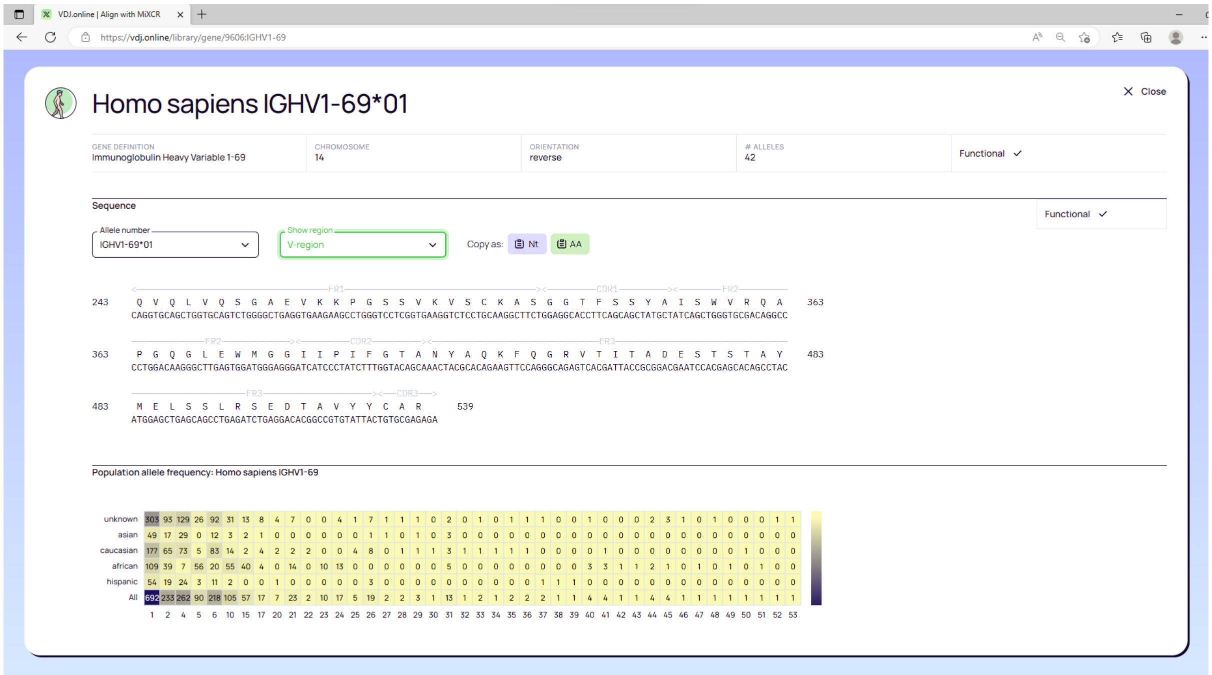


**Supplemental Figure S3.** VDJ.online - The database of adaptive immunity genes and alleles for species and receptor types.

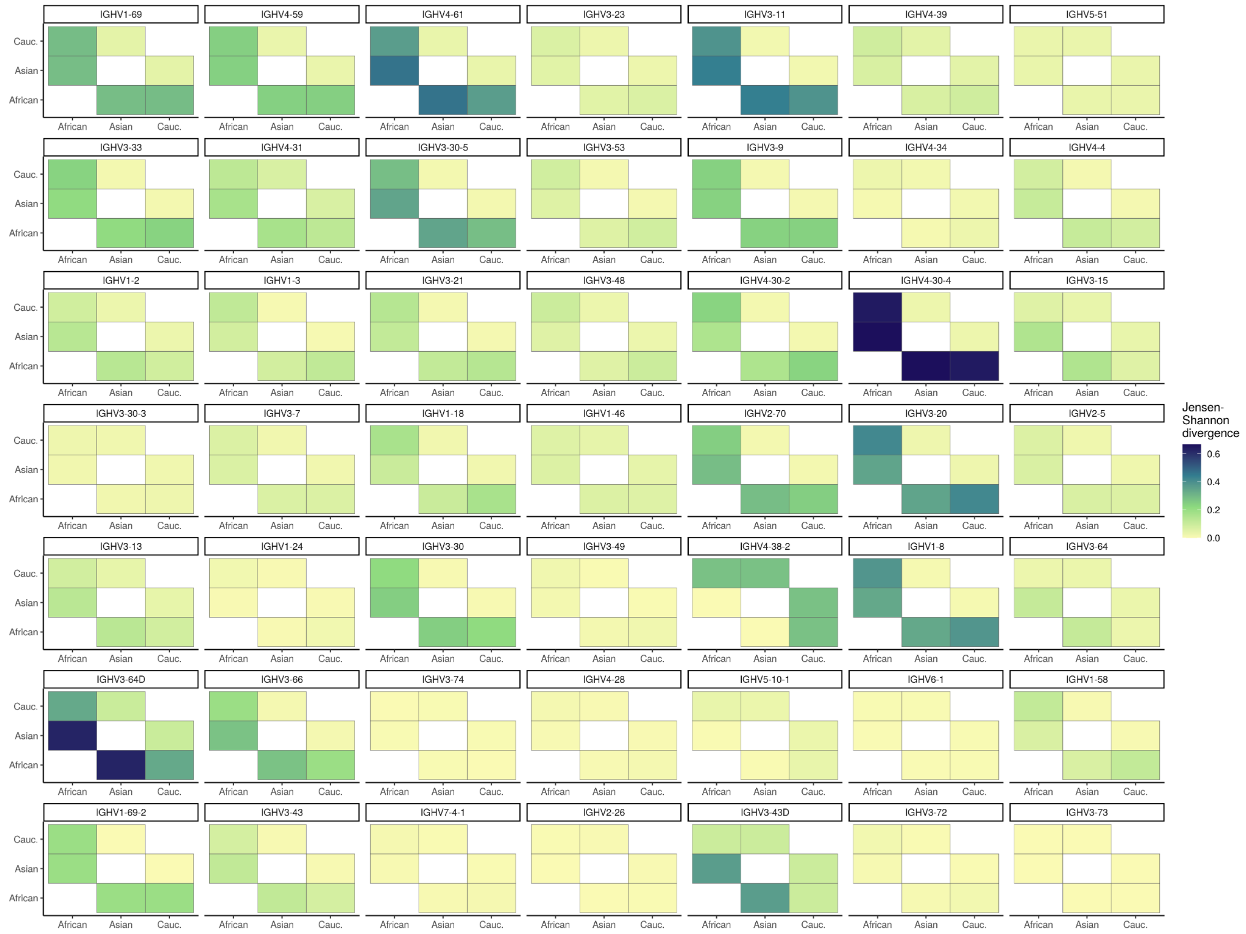


**Supplemental Figure S4.** Jensen-Shannon divergence of allele frequency distributions between major ethnic groups.

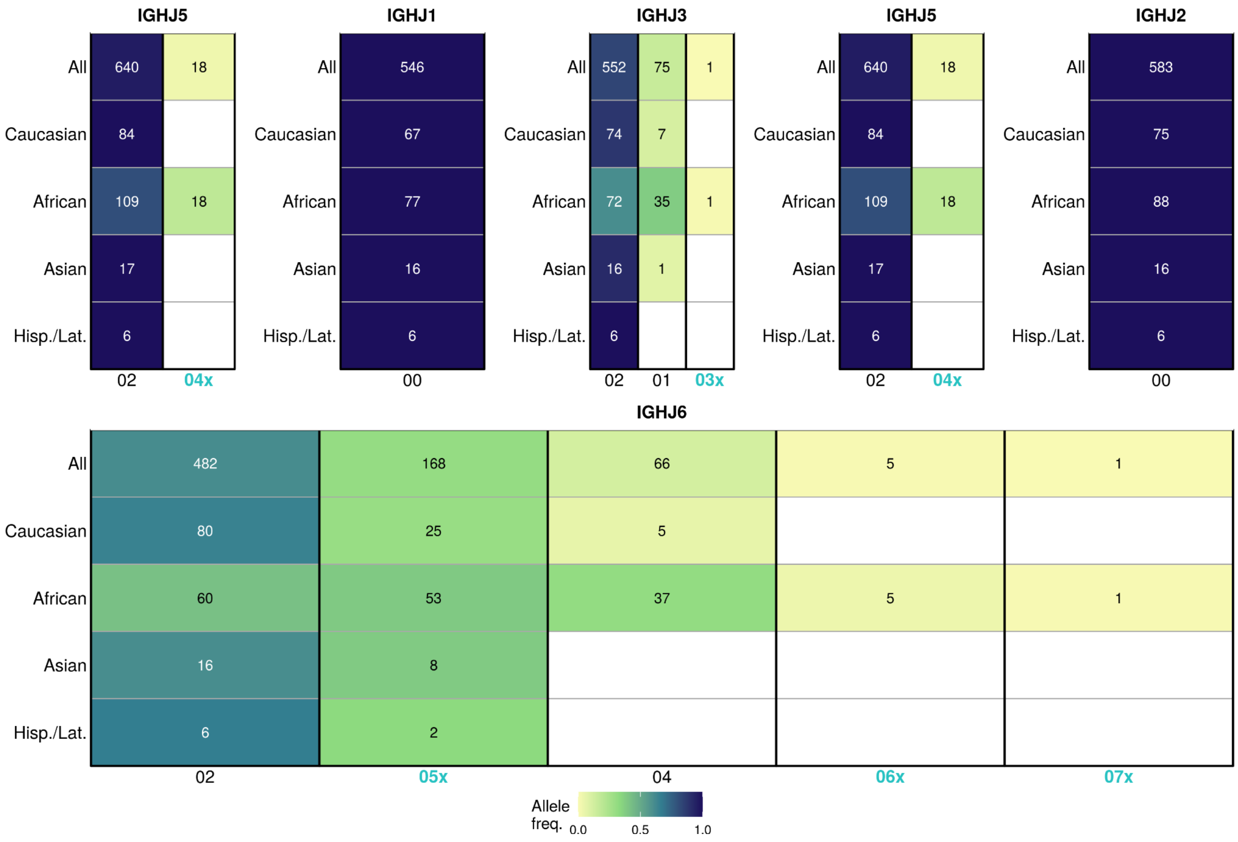


**Supplemental Figure S5. IGHJ gene allele frequencies in major ethnic groups.** Each column in heatmaps represents a particular allele, each row - major ethnic group. Numbers for novel alleles are colored in green, and concatenated with letter ‘x’; alleles are ordered by allele frequency in the general population. Color represents the allele frequency within ethnic groups; numbers in cells represent the number of occurrences of the corresponding allele.

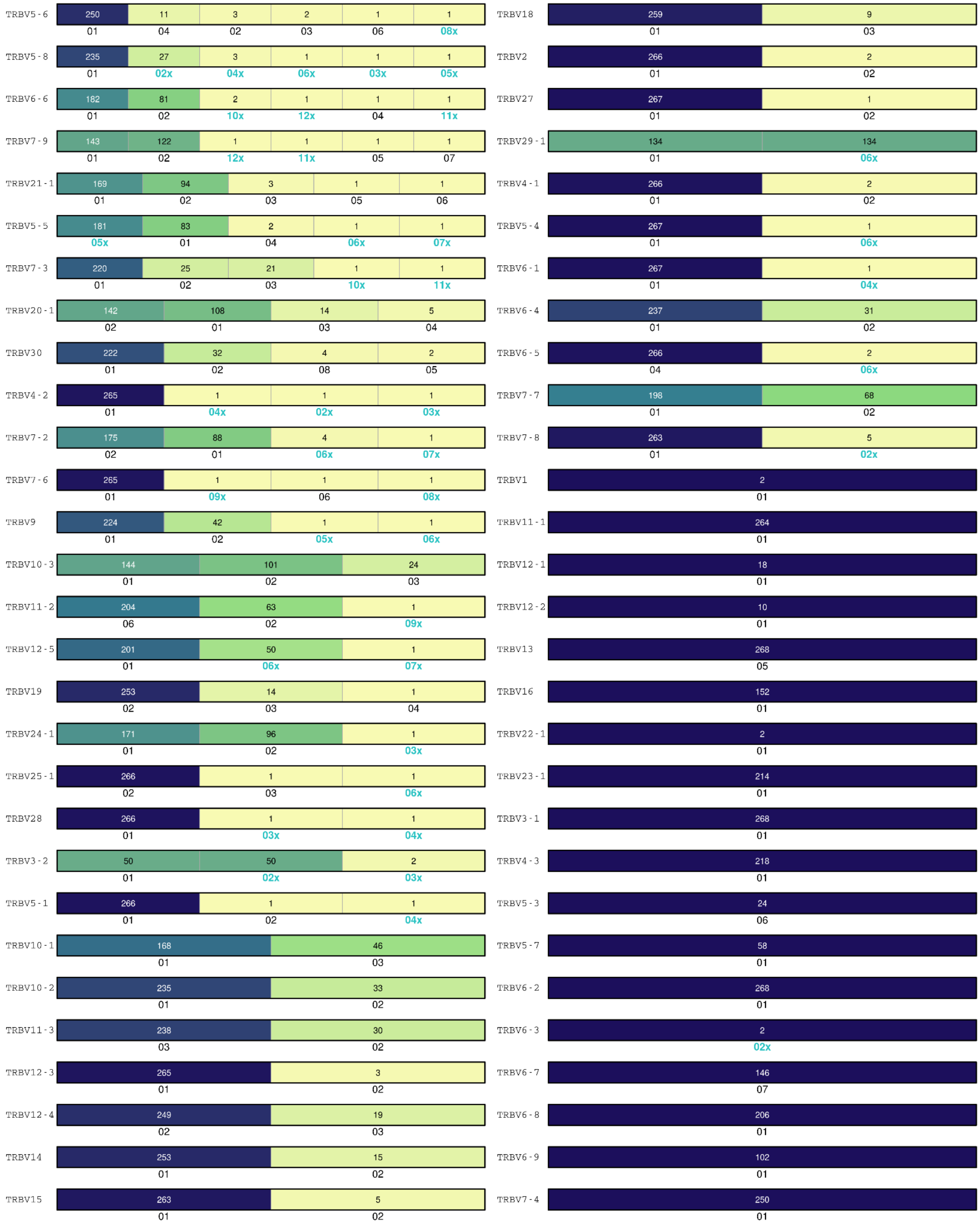

TRBV5-6
250 11 3 2 1 1
01 04 02 03 06 08x
TRBV5-8
235 27 3 1 1 1
01 02x 04x 06x 03x 05x
TRBV6-6
182 81 2 1 1 1
01 02 10x 12x 04 11x
TRBV7-9
143 122 1 1 1 1
01 02 12x 11x 05 07
TRBV21-1
169 94 3 1 1
01 02 03 05 06
TRBV5-5
181 83 2 1 1
05x 01 04 06x 07x
TRBV7-3
220 25 21 1 1
01 02 03 10x 11x
TRBV20-1
142 108 14 5
02 01 03 04
TRBV30
222 32 4 2
01 02 08 05
TRBV4-2
265 1 1 1
01 04x 02x 03x
TRBV7-2
175 88 4 1
02 01 06x 07x
TRBV7-6
265 1 1 1
01 09x 06 08x
TRBV9
224 42 1 1
01 02 05x 06x
TRBV10-3
144 101 24
01 02 03
TRBV11-2
204 63 1
06 02 09x
TRBV12-5
201 50 1
01 06x 07x
TRBV19
253 14 1
02 03 04
TRBV24-1
171 96 1
01 02 03x
TRBV25-1
266 1 1
02 03 06x
TRBV28
266 1 1
01 03x 04x
TRBV3-2
50 50 2
01 02x 03x
TRBV5-1
266 1 1
01 02 04x
TRBV10-1
168 46
01 03
TRBV10-2
235 33
01 02
TRBV11-3
238 30
03 02
TRBV12-3
265 3
01 02
TRBV12-4
249 19
02 03
TRBV14
253 15
01 02
TRBV15
263 5
01 02
TRBV18
259 9
01 03
TRBV2
266 2
01 02
TRBV27
267 1
01 02
TRBV29-1
134 134
01 06x
TRBV4-1
266 2
01 02
TRBV5-4
267 1
01 06x
TRBV6-1
267 1
01 04x
TRBV6-4
237 31
01 02
TRBV6-5
266 2
04 06x
TRBV7-7
198 68
01 02
TRBV7-8
263 5
01 02x
TRBV1
2
01
TRBV11-1
264
01
TRBV12-1
18
01
TRBV12-2
10
01
TRBV13
268
05
TRBV16
152
01
TRBV22-1
2
01
TRBV23-1
214
01
TRBV3-1
268
01
TRBV4-3
218
01
TRBV5-3
24
06
TRBV5-7
58
01
TRBV6-2
268
01
TRBV6-3
2
02x
TRBV6-7
146
07
TRBV6-8
206
01
TRBV6-9
102
01
TRBV7-4
250
01

**Supplemental Figure S6. TRBV gene allele  frequencies.** Each column in heatmaps represents a particular allele. Numbers for novel alleles are colored in green, and concatenated with letter ‘x’; alleles are ordered by allele frequency in the general population. Color represents the allele frequency within the general population; numbers in cells represent the number occurrences of the corresponding allele.

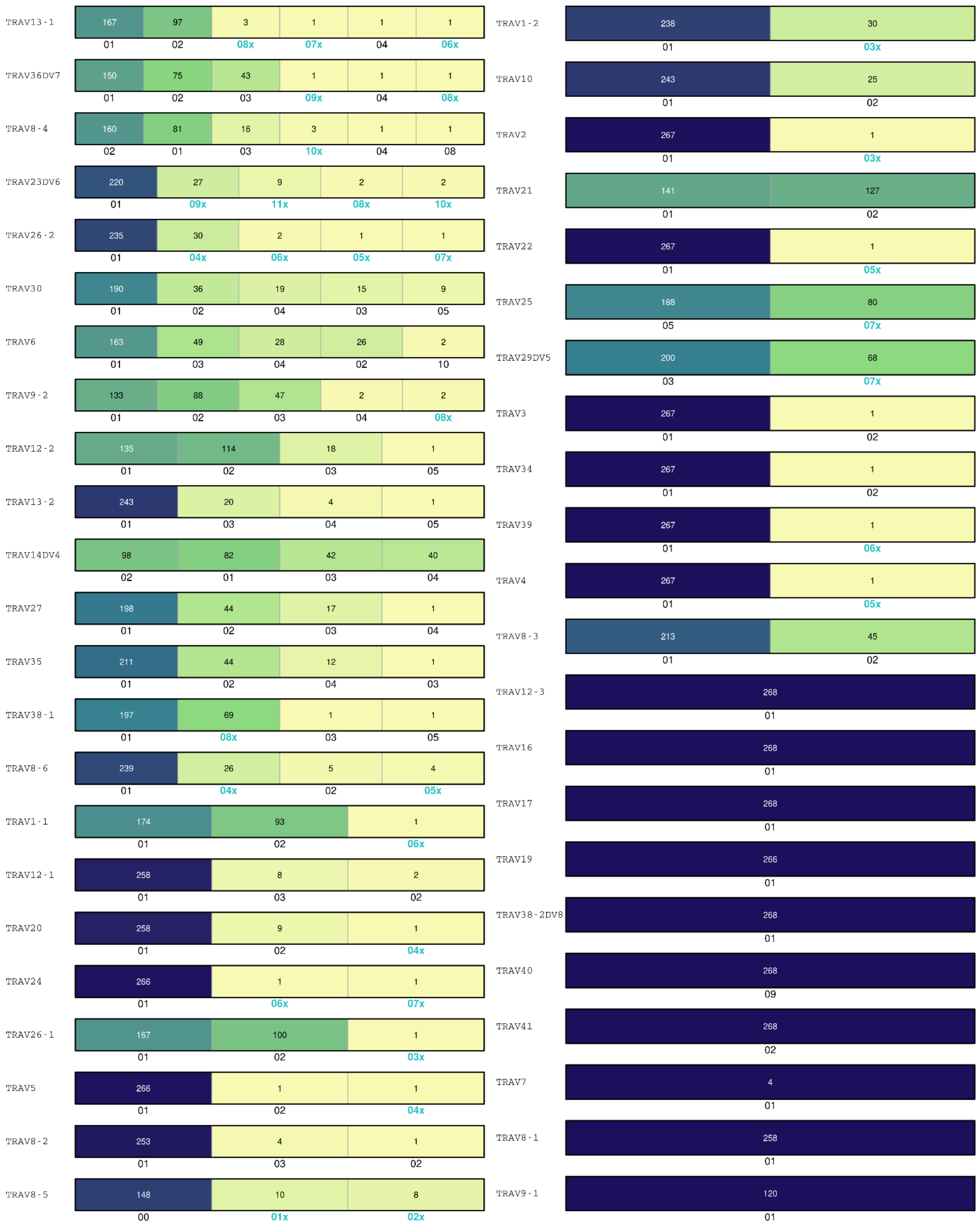
TRAV13-1
167 97 3 1 1 1
01 02 08x 07x 04 06x
TRAV36DV7
150 75 43 1 1 1
01 02 03 09x 04 08x
TRAV8-4
160 81 16 3 1 1
02 01 03 10x 04 08
TRAV23DV6
220 27 9 2 2
01 09x 11x 08x 10x
TRAV26-2
235 30 2 1 1
01 04x 06x 05x 07x
TRAV30
190 36 19 15 9
01 02 04 03 05
TRAV6
163 49 28 26 2
01 03 04 02 10
TRAV9-2
133 88 47 2 2
01 02 03 04 08x
TRAV12-2
135 114 18 1
01 02 03 05
TRAV13-2
243 20 4 1
01 03 04 05
TRAV14DV4
98 82 42 40
02 01 03 04
TRAV27
198 44 17 1
01 02 03 04
TRAV35
211 44 12 1
01 02 04 03
TRAV38-1
197 69 1 1
01 08x 03 05
TRAV8-6
239 26 5 4
01 04x 02 05x
TRAV1-1
174 93 1
01 02 06x
TRAV12-1
258 8 2
01 03 02
TRAV20
258 9 1
01 02 04x
TRAV24
266 1 1
01 06x 07x
TRAV26-1
167 100 1
01 02 03x
TRAV5
266 1 1
01 02 04x
TRAV8-2
253 4 1
01 03 02
TRAV8-5
148 10 8
00 01x 02x
TRAV1-2
238 30
01 03x
TRAV10
243 25
01 02
TRAV2
267 1
01 03x
TRAV21
141 127
01 02
TRAV22
267 1
01 05x
TRAV25
188 80
05 07x
TRAV29DV5
200 68
03 07x
TRAV3
267 1
01 02
TRAV34
267 1
01 02
TRAV39
267 1
01 06x
TRAV4
267 1
01 05x
TRAV8-3
213 45
01 02
TRAV12-3
268
01
TRAV16
268
01
TRAV17
268
01
TRAV19
266
01
TRAV38-2DV8
268
01
TRAV40
268
09
TRAV41
268
02
TRAV7
4
01
TRAV8-1
258
01
TRAV9-1
120
01

**Supplemental Figure S7. TRAV gene allele  frequencies.** Each column in heatmaps represents a particular allele. Numbers for novel alleles are colored in green, and concatenated with letter ‘x’; alleles are ordered by allele frequency in the general population. Color represents the allele frequency within the general population; numbers in cells represent the number of occurrences of the corresponding allele.
